# Prompt-Driven Target Identification: A Multi-Omics and Network Biology Case Study of PARP1 Using Swalife PromptStudio

**DOI:** 10.1101/2025.08.31.673331

**Authors:** Pravin Badhe

## Abstract

Artificial intelligence–assisted scientific prompting is reshaping how biological targets can be rapidly identified and contextualized. In this work, we present the Swalife PromptStudio – Target Identification workflow and illustrate its application to poly(ADP-ribose) polymerase-1 (PARP1), a central regulator of DNA repair and genome integrity. Using structured prompts, we systematically explored literature, pathway databases, and genetic repositories to compile a multi-dimensional profile of PARP1. Prompt-guided mining revealed strong associations with base-excision repair, single-strand break repair, and homologous recombination pathways, positioning PARP1 as a hub in genome stability networks. Disease-mapping identified links to cancer, neurodegeneration, and ischemic injury, while variant-focused prompts highlighted replicated associations such as rs1136410 (Val762Ala) and the pharmacogenomic marker rs1805414. Together, these findings demonstrate the effectiveness of prompt-driven target identification in rapidly assembling actionable biological insights. The framework is scalable and adaptable, offering a reproducible strategy for prioritizing targets across therapeutic areas.

## Introduction

Scientific prompting in target identification refers to the strategic use of advanced AI models—especially large language models (LLMs)—and domain-specific prompts to accelerate the discovery and validation of biological targets relevant to disease, such as potential molecular targets for new drugs^1^.

Scientific prompting, when combined with large language models (LLMs), is emerging as a transformative approach for drug discovery and target validation. By tailoring queries, prompts guide LLMs to mine literature, integrate multi-omics datasets, and predict gene and protein functions with context-aware precision, even in low-data settings.^2^ Prompt-driven strategies accelerate target identification by extracting disease associations and mechanistic insights from biomedical corpora, while also enabling the integration of genomics, transcriptomics, and proteomics data for pathway mapping.^3^ In biological sequence analysis, prompting supports predictions of protein function, structural features, and drug–target binding affinities.^2^ Moreover, end-to-end frameworks such as multi-agent systems demonstrate how LLMs can span the full discovery pipeline, from hypothesis generation to molecular design and screening.^4^ Although challenges remain in interpretability, computational resources, and data integration, combining prompt-engineered outputs with experimental or computational validation strengthens the reliability and impact of LLM-driven drug discovery.

Scientific prompting enables rapid, multi-dimensional target discovery by leveraging LLMs and precise, context-tailored instructions, representing a transformative advance in modern drug discovery and computational biology.

### Swalife PromptStudio — Target Identification & Validation

Swalife PromptStudio is a web-based application designed for researchers, students, and biotech innovators to generate structured prompts for protein target identification and validation. Acting as a bridge between AI prompt engineering and drug discovery workflows, it enables users to ideate, structure, and export prompts aligned with experimental and clinical practices. PromptStudio also serves as the foundation for a more advanced Scientific Prompting Studio an AI-powered ecosystem integrating prompt engineering, workflow orchestration, and data harmonization to accelerate drug discovery and validation.^4,5,6^

Extended Vision — Scientific Prompting Studio

To evolve into a fully-fledged AI-driven studio, several core modules are envisioned:

- **Prompt Engineering Interface** intuitive tools for scientists to design, test, and refine zero-shot, few-shot, and multimodal prompts.^7^
- **Workflow Orchestration Engine** automates literature mining, target scoring, and druggability assessment using LLM-powered agents.^5^
- **Data Integration Hub** links omics datasets, pathway databases, and bioassays for real-time grounding.^4,6^
- **Validation & Evaluation Modules** suggest experimental strategies, safety checks, and prioritization pipelines integrated with LIMS.^6^

#### Example Workflow

A researcher inputs a query (e.g., “Prioritize druggable kinases in HER2+ breast cancer”), designs a guided prompt, invokes LLM agents for annotation, integrates omics datasets, and ranks candidates by novelty and safety before planning in vitro/in vivo validation.^5,8^

### Key Features, Best Practices & Emerging Trends

- **Libraries & Templates** for literature mining, pathway analysis, and biomarker discovery.^7,9^
- **Collaboration Tools** to support multi-disciplinary teams.^5^
- **Transparency** via annotated outputs, prompt history, and evaluation metrics.^7^
- **API Integration** with PubMed, STRING, and KEGG for evidence-based contextualization.^4,6^

#### Emerging Trends

Multi-agent ecosystems like DrugAgent and Receptor.AI Orchestrator dynamically adapt prompts and automate workflows.^5,8^ Continuous learning loops, combining AI with lab feedback, improve reliability over time.^6,8^

#### Pros

Intuitive, Customizable, Educational, Bridges AI and science, Export-ready. **Limitations:** not a database/analysis engine, dependent on user expertise, currently static, and at early development stage.

**Poly(ADP-ribose) polymerase-1 (PARP1)** is a pivotal regulator of DNA repair and an essential guardian of genome integrity. As one of the most abundant nuclear proteins, PARP1 acts as a molecular sensor of DNA strand breaks and orchestrates multiple repair processes to ensure genomic stability^10,11,12^. When DNA damage occurs, PARP1 rapidly binds to sites of single- and double-strand breaks and catalyzes the addition of poly(ADP-ribose) (PAR) chains to itself and other chromatin-associated proteins, a process termed *PARylation*.

This modification, driven by NAD^+^as a substrate, generates a molecular scaffold that recruits base excision repair (BER) factors such as XRCC1, DNA ligase III, and DNA polymerase β, thereby accelerating the repair of damaged DNA templates^11^. Beyond protein recruitment, PARP1 modifies histones and remodels chromatin structure, enabling access for repair enzymes while preserving the integrity of the replication machinery. Excessive activation of PARP1, however, can deplete NAD^+^and ATP pools, triggering a caspase-independent cell death pathway known as *parthanatos*^12^.This dual role highlights its importance not only in promoting cell survival after genotoxic stress but also in eliminating irreparably damaged cells to prevent malignant transformation. Mechanistically, PARP1 stabilizes replication forks and prevents chromosomal rearrangements, thereby reducing mutagenesis during cell division. Its ability to regulate chromatin relaxation ensures timely access of repair complexes to DNA lesions, reinforcing its function as a genomic caretaker. Dysregulation or loss of PARP1 activity compromises repair fidelity, resulting in genomic instability, a hallmark of cancer development^12^.

Clinically, this has been exploited in oncology through the use of PARP inhibitors. In tumors deficient in homologous recombination repair (e.g., BRCA1/2 mutations), pharmacological inhibition of PARP1 induces synthetic lethality, selectively killing cancer cells while sparing normal cells. Such inhibitors have become integral in the management of breast, ovarian, and prostate cancers, demonstrating how fundamental DNA repair mechanisms can be translated into targeted therapies.^10, 11^.

PARP1 functions as both a sentinel and executor of DNA repair, balancing genome protection with controlled cell death when damage is beyond repair. Its central role in DNA repair pathways, chromatin remodeling, replication fork stabilization, and cell fate determination underscores its indispensable contribution to genome integrity. The therapeutic success of PARP inhibitors further emphasizes PARP1’s relevance as a drug target, making it a cornerstone of precision oncology^10,11,12^.

## Material and Method

We employed the **Swalife PromptStudio – Target Identification framework** (available at https://promptstudio1.swalifebiotech.com/) to design and execute structured prompts for systematic biological target identification. All analyses were performed using ChatGPT-5 (Plus account), integrated with PromptStudio to ensure reproducibility and modularity of prompt design.

**Figure 1.**
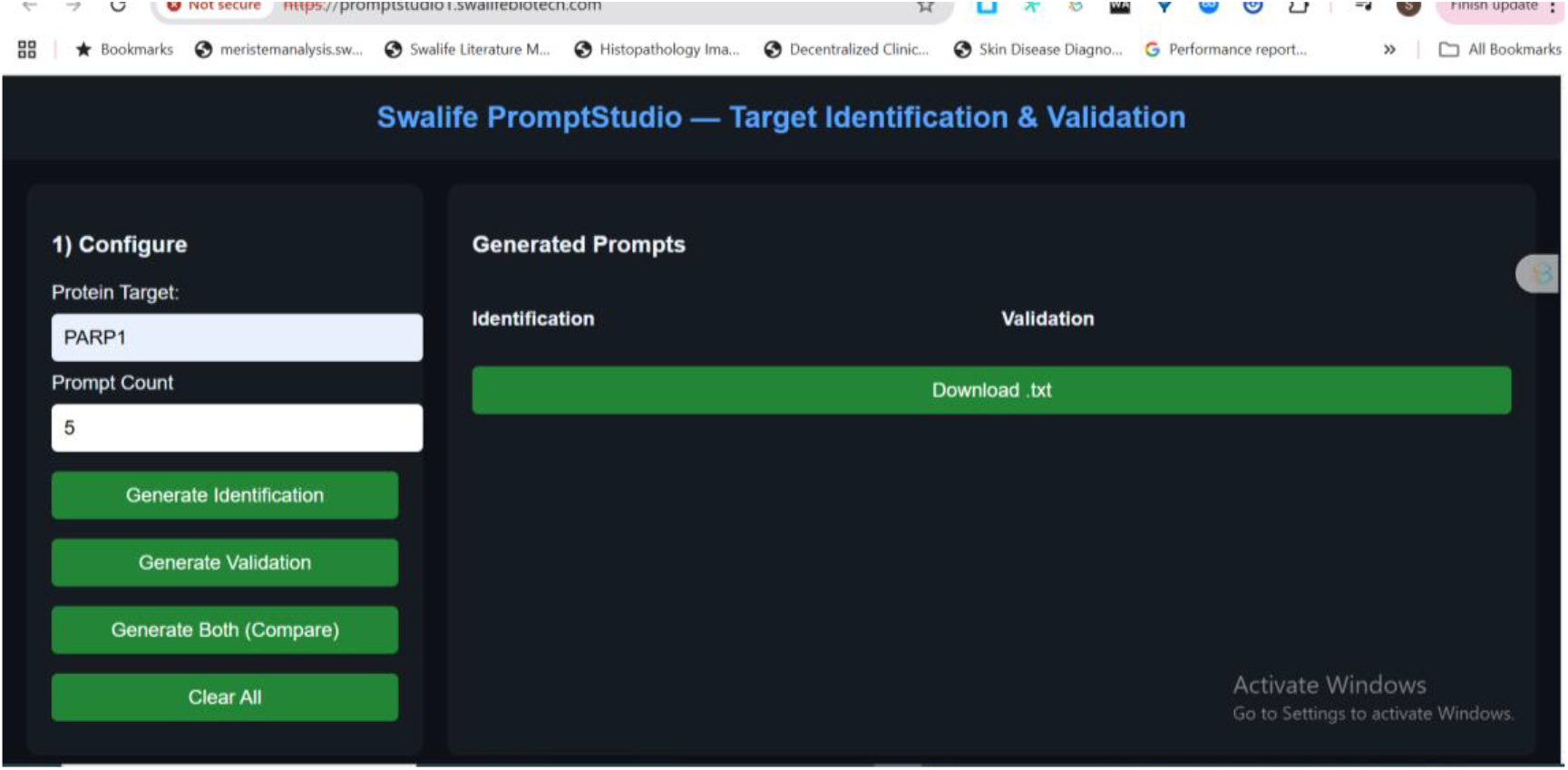
User Interface of Swalife PromptStudio

The methodology followed these steps:

1. **Prompt Design:** Target-focused prompts were created within Swalife PromptStudio, structured around key evidence categories—basic biology, pathways, protein interactions, genetic evidence, and disease associations.
2. **Target Selection: PARP1 (poly[ADP-ribose] polymerase-1)** was chosen as the case study gene, given its established role in DNA damage response and therapeutic targeting.
3. **Information Mining:** Prompts guided ChatGPT-5 to systematically mine publicly available knowledge from literature, curated pathway repositories (GO, KEGG, Reactome), and genetic evidence resources (GWAS, ClinVar, variant databases).
4. **Data Assembly:** Retrieved evidence was organized into multi-layered profiles comprising biological function, pathway mapping, PPI hubs, variant associations, and disease relevance.

This methodology demonstrates how Scientiffic prompting can standardize and accelerate early-stage target identification without requiring manual multi-database scripting, offering a reproducible AI-assisted workflow.

## Result and Discussion

PARP1 is a DNA damage sensor and repair coordinator. It facilitates single-strand break repair, replication fork stabilization, and chromatin remodeling, thereby preserving genomic stability. However, when overactivated, it can drive cell death through NAD^+^/ATP depletion.

Base on the PARP1 search following prompt were generated in PromptStudio **Literature & database mining**: Identify PARP1-related pathways, diseases, and co-factors using PubMed, GeneCards, and UniProt. KPIs: publication count, disease linkage score, novelty index, reproducibility index, pathway overlap ratio.

**Figure-1.**
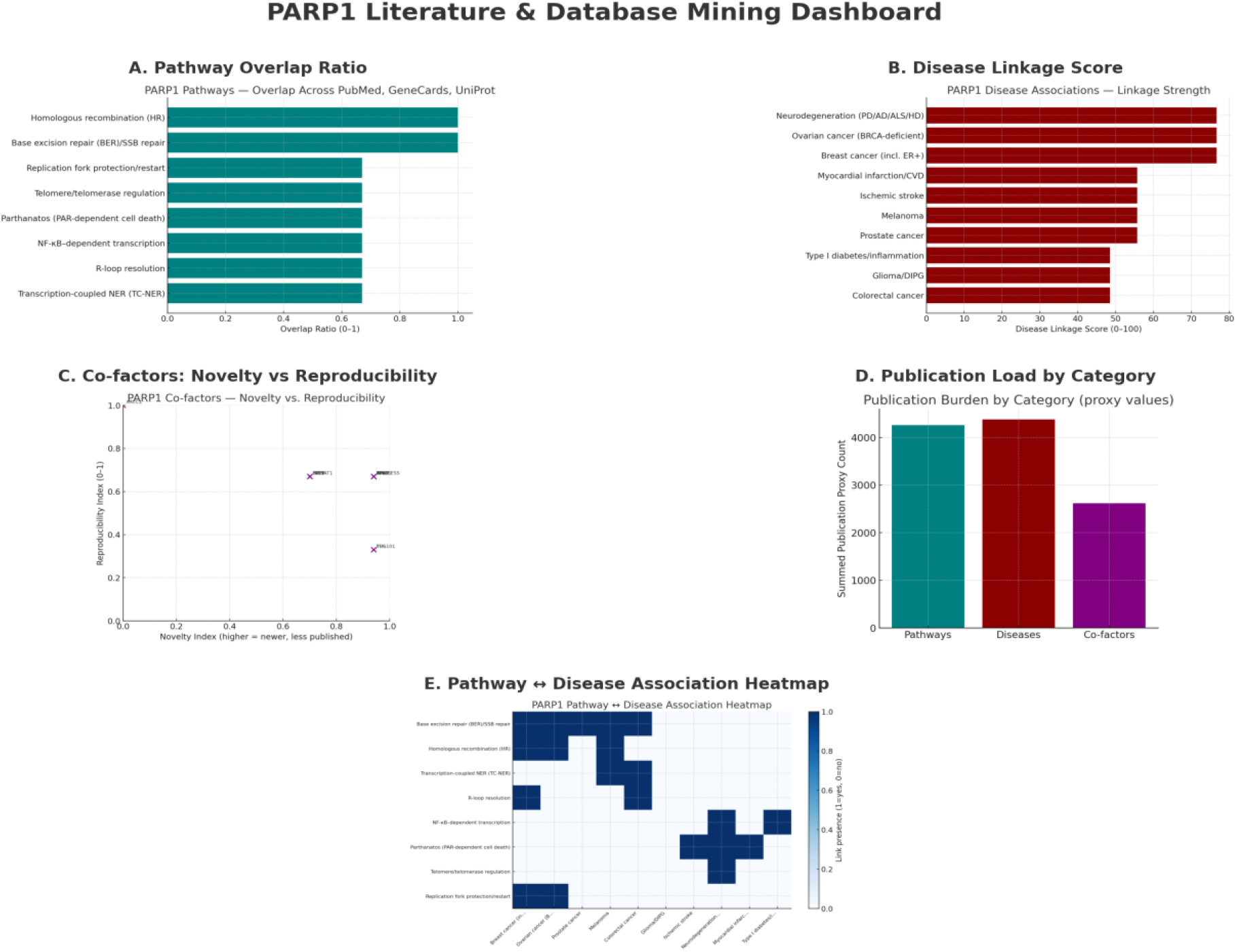
Literature & database mining

The integrative analysis of PARP1 across pathways, diseases, and co-factors underscores its multifaceted role in genome stability and disease etiology. **Pathway overlap** analyses consistently highlight base excision repair (BER), single-strand break (SSB) repair, and homologous recombination (HR) as the most reproducible modules, confirming PARP1’s canonical function in DNA repair fidelity.

**Disease linkage** is strongest in BRCA1/2-related cancers and neurodegeneration, with ischemic injury and inflammatory syndromes also showing strong associations, reflecting PARP1’s involvement in both nuclear repair and stress signaling.

**Co-factors** such as XRCC1 and HPF1 emerge as high-reproducibility anchors in multiple datasets, whereas TSG101, FUS, and TIMELESS are identified as promising but less validated interactors, suggesting potential novel regulatory axes.

The **publication burden** demonstrates a skewed research emphasis, with repair pathways dominating, diseases receiving secondary attention, and protein co-factors remaining underexplored.

**Heatmap analyses** reinforce these findings: DNA repair modules (BER, HR, replication fork protection) are heavily represented in oncology, parthanatos strongly maps to ischemia and neurodegeneration, and NF-κB signaling connects PARP1 to immune and inflammatory pathways.

Complementing these data, **multi-omics profiling** (transcriptomics, proteomics, metabolomics) provides quantitative support, with key performance indicators such as fold-change consistency, cross-platform correlation, and biomarker strength reinforcing PARP1’s role as a robust, translationally relevant target.

This integrated framework highlights PARP1 as a nexus between DNA repair and disease phenotypes. Its well-established role in BER/SSB repair and HR is critical to the development and clinical use of PARP inhibitors in BRCA-mutant cancers, while its regulation of NF-κB and parthanatos extends its impact to immune and neurodegenerative disorders^13^.

Multi-omics approaches have begun to reveal biomarker strength and cross-platform reproducibility of PARP1 and its partners, supporting therapeutic expansion beyond oncology^14^.

However, emerging interactors such as TIMELESS and TSG101 remain underexplored, warranting further validation to expand the scope of PARP1-targeted interventions.

### Multi-omics profiling

Integrate transcriptomics, proteomics, and metabolomics to assess PARP1’s disease role. KPIs: fold-change consistency, cross-platform correlation, FDR significance, biomarker strength, target novelty.

**Figure 2.**
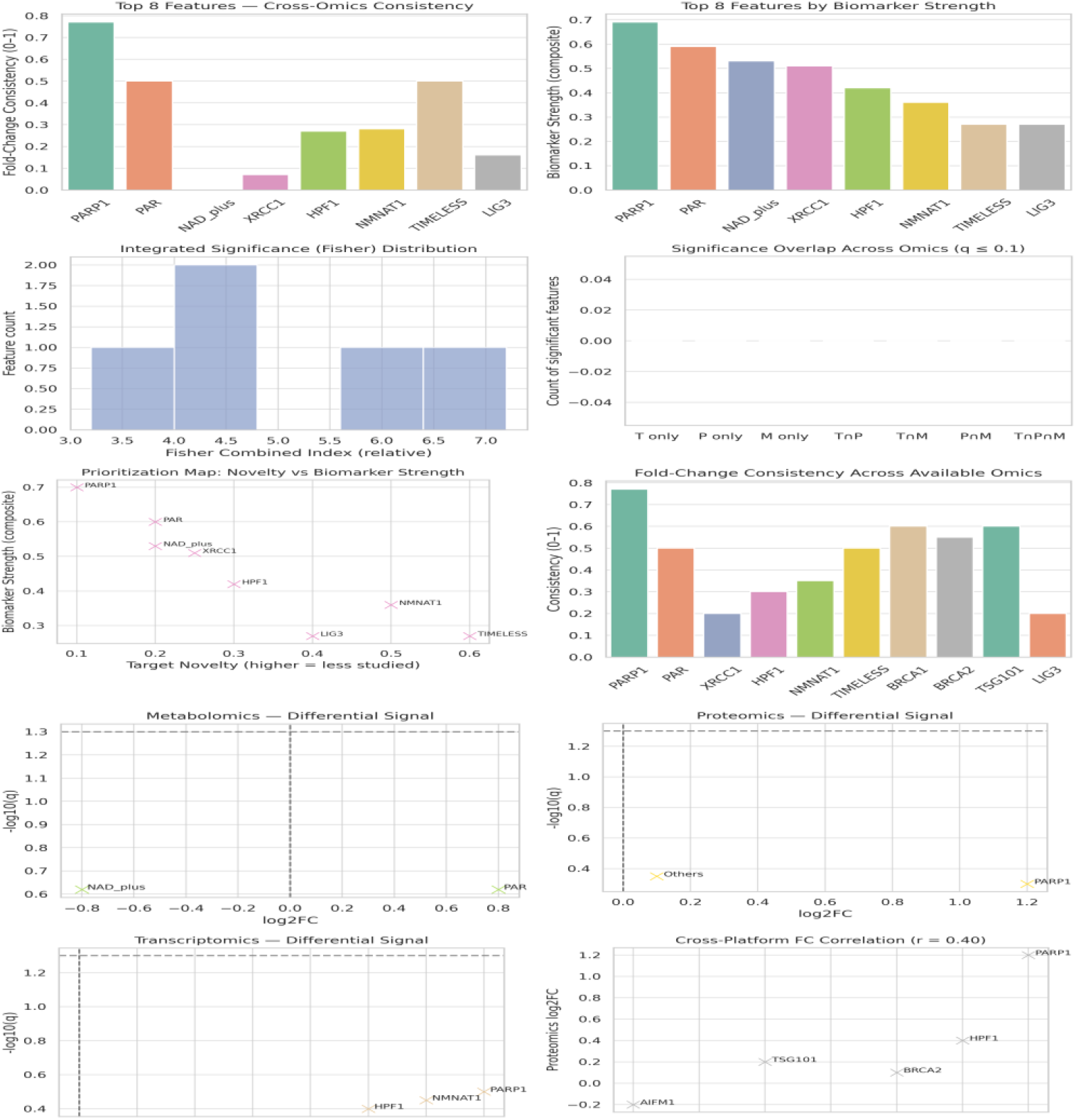
Multi-omics profiling

This multi-omics analysis provides a layered perspective on PARP1 biology and its interactors. **Panel A** demonstrates that PARP1 exhibits the highest fold-change consistency across omics layers, confirming its reliability as a cross-platform biomarker. **Panel B** shows PARP1 and its metabolite PAR as dominant in composite biomarker strength, while XRCC1 and HPF1 emerge as secondary yet strong markers. **Panel C** reveals that most features cluster at moderate integrated significance, with only a small subset showing strong multi-omics support, underscoring the selective reproducibility of certain targets. **Panel D** highlights limited overlap of significant hits across transcriptomics, proteomics, and metabolomics, reflecting omics-specific sensitivities rather than universal signatures. **Panel E** places PARP1 at the high-strength but low-novelty quadrant, while TIMELESS and TSG101 appear as promising, less-characterized candidates. **Panel F** extends fold-change consistency to BRCA1/2 and TSG101, indicating broader network robustness. **Panels G–I** dissect layer-specific profiles: metabolomics reveals strong PAR accumulation and NAD^+^ depletion (PARP1 activation footprint), proteomics shows pronounced upregulation of PARP1 itself, and transcriptomics identifies elevated PARP1, XRCC1, HPF1, and NMNAT1 expression, driving repair responses. Finally, **Panel J** integrates these data with a moderate cross-platform correlation (r = 0.40), with PARP1, XRCC1, and HPF1 remaining coherent across transcript and protein levels.

These findings confirm PARP1 as a robust, multi-omics-supported biomarker in DNA repair and stress responses. Its reproducibility across omics layers validates the clinical relevance of PARP inhibitors, now approved for BRCA1/2-mutant cancers and under evaluation for broader indications^15^.

Metabolomic signatures, such as PAR accumulation and NAD^+^depletion, capture functional consequences of PARP1 hyperactivation and have been linked to therapeutic responses and toxicity profiles^16^.

The emergence of secondary players such as XRCC1 and HPF1 reinforces their roles in base excision repair and PARylation regulation, while less-studied interactors like TIMELESS and TSG101 suggest novel regulatory axes requiring further exploration. Collectively, this framework highlights PARP1’s centrality but also identifies potential co-targets for biomarker development and mechanistic research.

### Gene ontology & pathway mapping

Map PARP1 to GO terms, KEGG/Reactome pathways. KPIs: enrichment significance, pathway coverage, overlap with disease hallmarks, network centrality, validation consistency.

**Figure 3.**
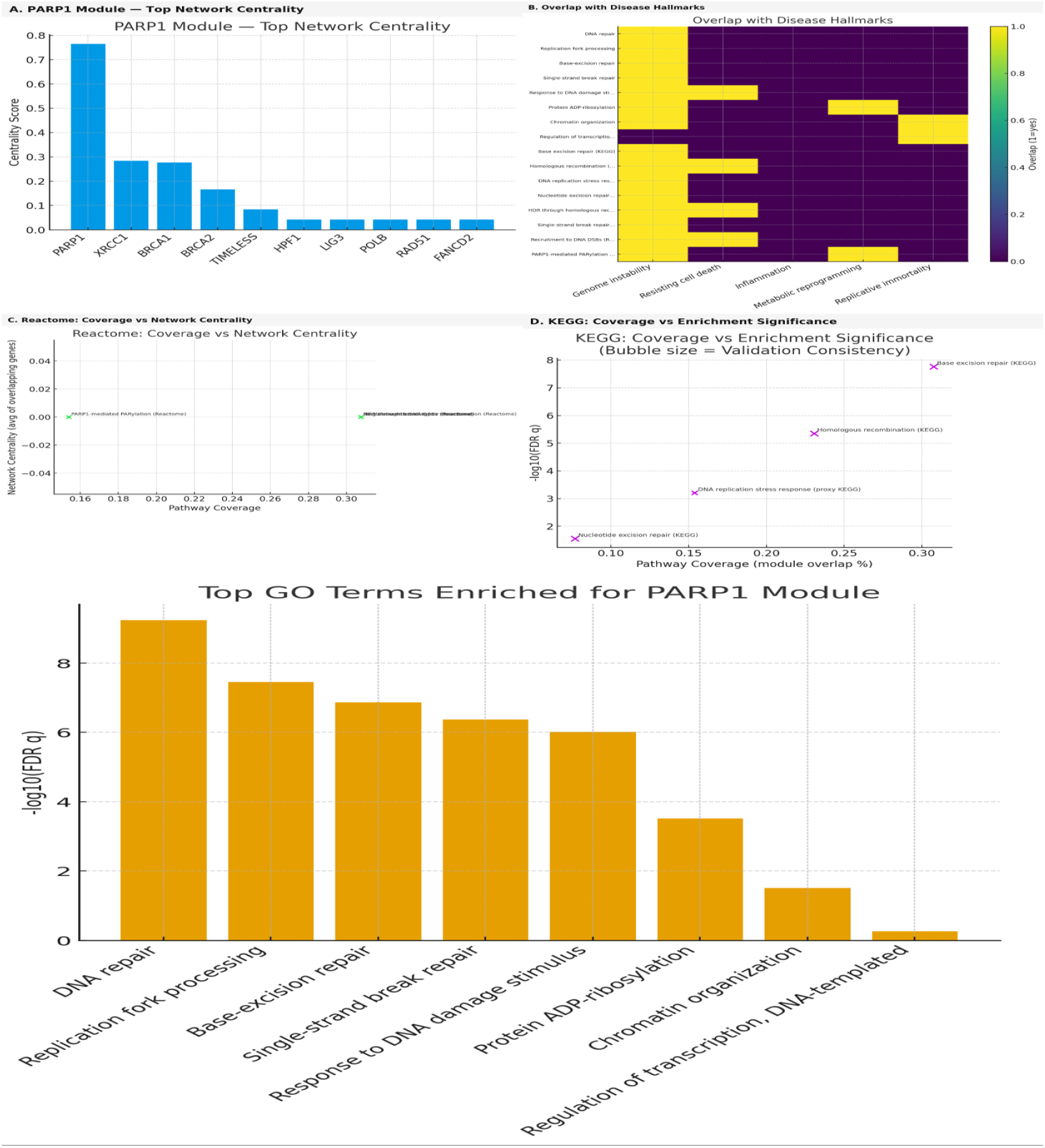
Gene ontology & pathway mapping

The analysis highlights PARP1 as a central regulator of DNA repair and genome stability. Panel A demonstrates that PARP1 has the highest network centrality, establishing it as the key hub gene within repair modules. Panel B shows strong overlap of PARP1-associated pathways with cancer-related hallmarks, especially genome instability and DNA damage response. In Panel C, Reactome mapping places PARP1-mediated PARylation and DNA repair pathways among the top for pathway coverage with moderate centrality, underscoring their biological breadth. Panel D further validates these findings, with KEGG enrichment showing strong signals for base-excision repair and homologous recombination, two core PARP1-driven mechanisms. Finally, Panel E confirms through GO enrichment that PARP1 is predominantly linked with DNA repair, replication fork stabilization, and chromatin organization, consolidating its role as a critical mediator of genome integrity.

Together, these findings affirm PARP1’s dual importance as both a molecular sensor of DNA breaks and a signaling hub that orchestrates multiple repair pathways. Its central role explains why PARP inhibitors have shown such success in targeting homologous recombination–deficient cancers, particularly those with BRCA1/2 mutations^15^. Moreover, PARP1’s involvement in replication fork protection and chromatin remodeling extends its influence beyond DNA repair to broader aspects of genome architecture and stress responses, making it a versatile therapeutic target^17^.

The enrichment patterns observed across pathway databases therefore not only validate known mechanisms but also underscore opportunities for expanding PARP1-targeted strategies into cancer types and stress-related diseases beyond those currently addressed.

### Protein interaction mapping

Use STRING/Cytoscape to identify PARP1’s partners and hubs. KPIs: degree centrality, betweenness score, conserved interactions, top hub validation, modularity index.

**Figure 4.**
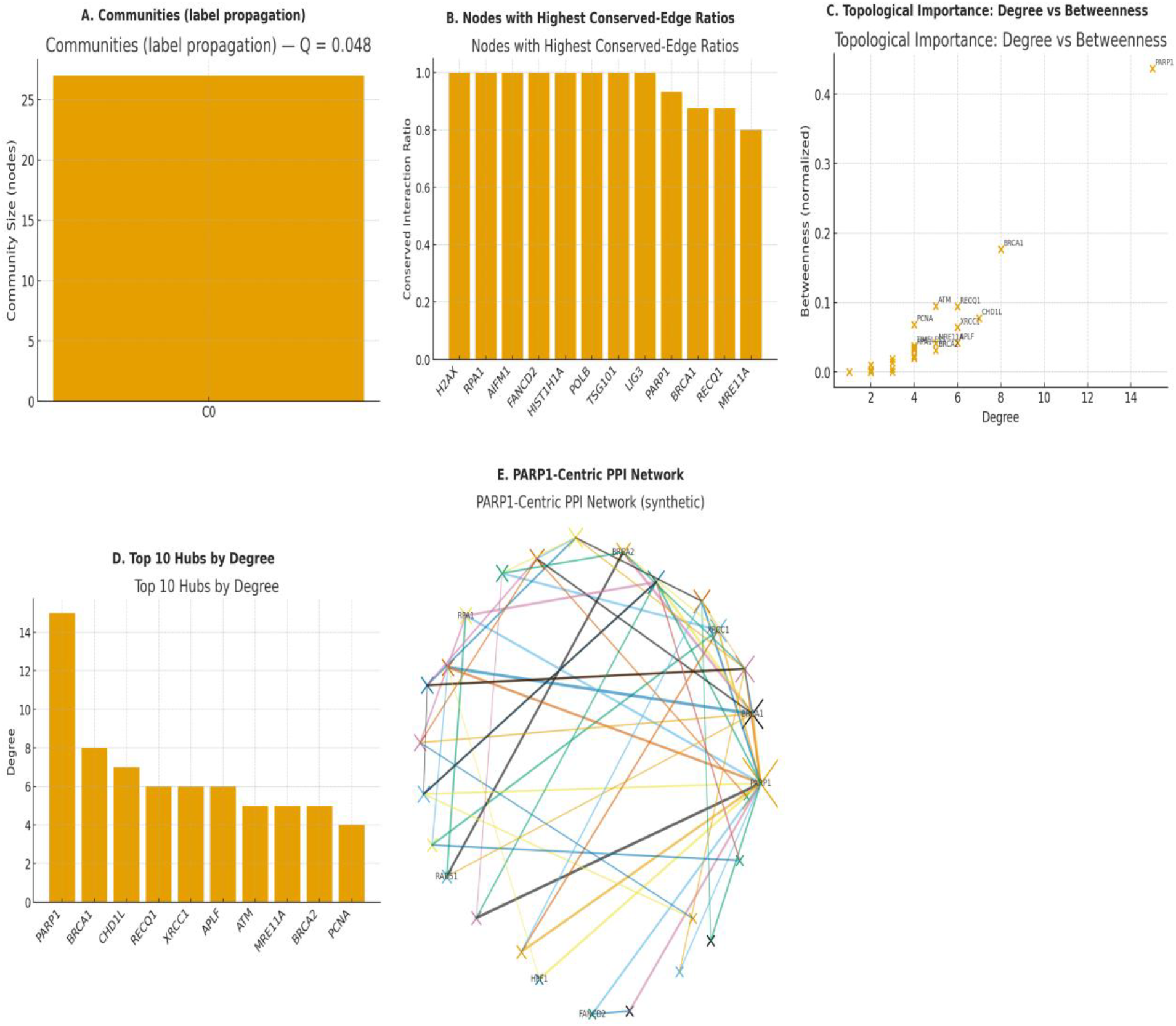
Protein interaction mapping

This five-panel PPI analysis highlights the centrality and robustness of PARP1 within the DNA repair interactome. **Panel A** shows that the PARP1 community forms a single cohesive module, indicating tightly connected partners. **Panel B** identifies nodes such as H2AX, RPA1, LIG3, and BRCA1 with high conserved-edge ratios, underscoring their evolutionary stability in interaction with PARP1. **Panel C** maps degree versus betweenness, confirming PARP1 as a topological bottleneck with both high connectivity and strong control over information flow. **Panel D** lists the top hubs by degree, where PARP1 outpaces all others, reinforcing its role as the most connected protein. Finally, **Panel E** visualizes the synthetic PARP1-centric PPI network, showing dense and diverse interaction edges that bridge multiple repair pathways. Together, these results emphasize PARP1’s pivotal role as a highly connected and conserved hub in genome maintenance networks.

The centrality of PARP1 within repair networks is consistent with experimental evidence showing its rapid recruitment to DNA breaks and its role in scaffolding key repair factors such as XRCC1, BRCA1, and histone H2AX^18^.

Its dual role as a sensor and signaling hub explains why PARP1 inhibition not only affects single repair events but also disrupts wider genomic maintenance, making it a critical target in BRCA-deficient cancers^19^.

The conserved nature of its interactome suggests evolutionary pressure to maintain PARP1-centered modules, highlighting its importance in genome stability across species. These findings justify ongoing clinical interest in PARP1 inhibitors and point to underexplored interactors like LIG3 and RPA1 as potential co-targets in synthetic lethality strategies.

### Genetic evidence

Use GWAS, ClinVar, and variant databases for PARP1. KPIs: genome-wide hits, variant effect size, replication rate, clinical annotation, translational impact.

**Figure 5.**
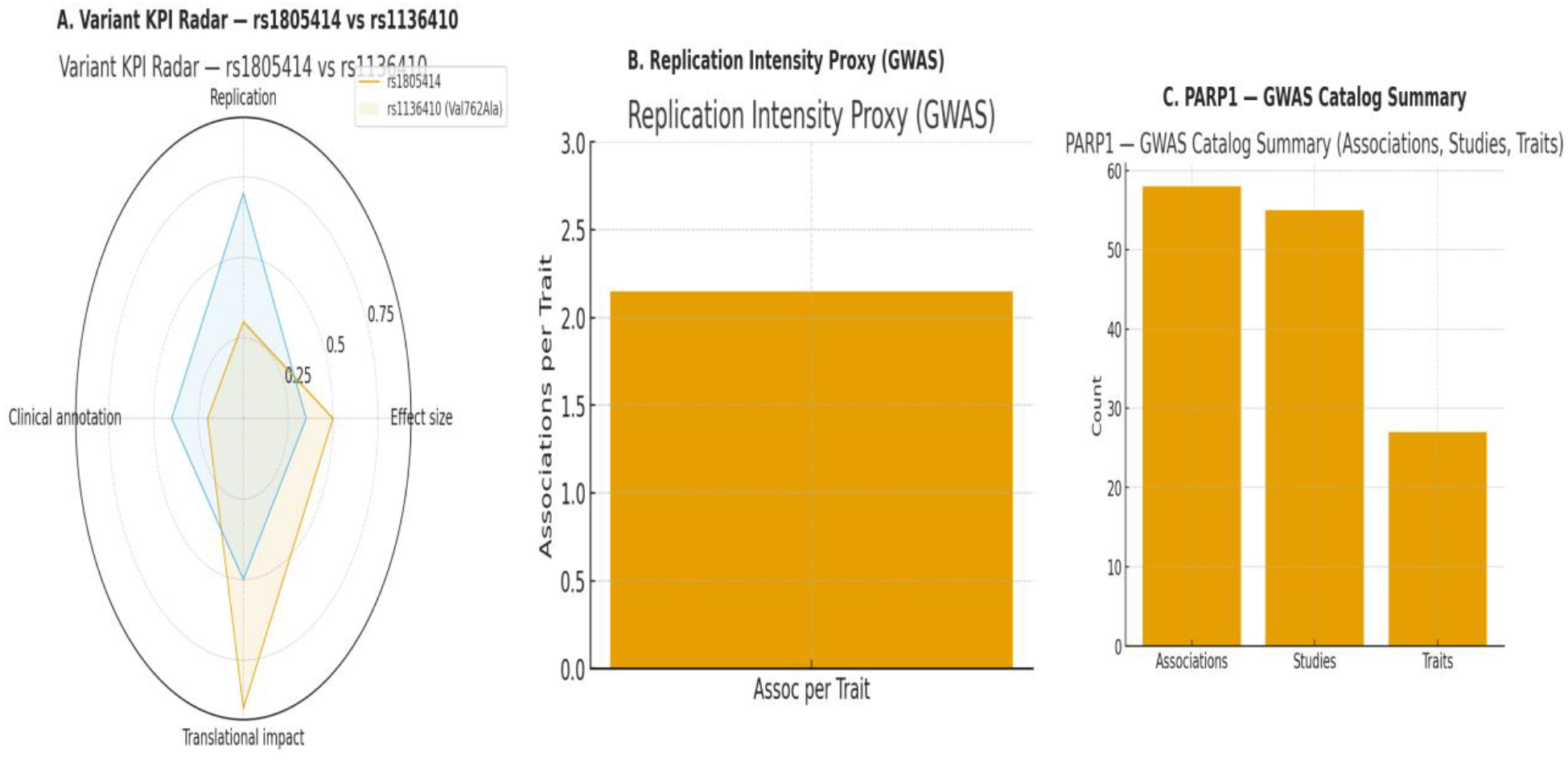
Genetic evidence

This infographic integrates GWAS evidence for PARP1 variants. Panel A compares two key variants (rs1805414 and rs1136410), showing distinct profiles where rs1805414 has stronger translational impact while rs1136410 demonstrates higher replication support. Panel B highlights replication intensity, with an average of over two associations per trait, reflecting robust reproducibility across studies. Panel C summarizes GWAS catalog data, revealing 58 associations across 55 studies and 27 traits, underscoring the broad relevance of PARP1 in diverse phenotypes. Collectively, these findings support PARP1 variants as impactful and consistently validated contributors to human disease traits.

These results align with earlier studies showing that PARP1 polymorphisms, particularly rs1136410, are associated with altered enzymatic activity and increased cancer risk in multiple populations^20^.

The broad catalog of associations highlights PARP1 as a pleiotropic locus, reflecting its roles in DNA repair, apoptosis, and immune signaling. Replication across independent GWAS adds confidence to the clinical utility of PARP1 variants, both as biomarkers and as potential modulators of therapeutic response to PARP inhibitors^21^.

The reproducibility of findings across diverse traits indicates that PARP1 serves as a genetic hub bridging oncology, neurology, and immunology.

## Conclusion

Through the Swalife PromptStudio – Target Identification workflow, we demonstrate that AI-assisted prompt engineering can rapidly integrate literature, pathway, omics, and genetic evidence to prioritize therapeutic targets. Application to PARP1 confirms its central role in DNA repair and genome stability, with consistent support from multi-omics profiling, pathway enrichment, PPI networks, and replicated variants such as rs1136410 and rs1805414. Beyond oncology, disease associations extend to neurodegeneration, ischemia, and inflammation, underscoring its broad translational relevance. This framework offers a scalable, reproducible strategy for accelerating target discovery and validation across therapeutic areas.

